# Mind the corner: Fillets in cryo-FIB lamella preparation to minimise sample loss caused by stress concentration and lamella breakage

**DOI:** 10.1101/2025.04.21.649880

**Authors:** Sergey Gorelick, Sailakshmi Velamoor, Patrick Cleeve, Sylvain Trépout, Le Ying, Vivek Naranbhai, Georg Ramm

## Abstract

Cryo-FIB milling of biological specimens is a critical and limiting step in the cryo-electron tomography workflow. Preparing electron-transparent cryo-lamellae is a serial, low-throughput process. Even with automation, a skilled operator can typically only produce 15–25 lamellae in a single cryo-FIB session. During sample handling, milling and transfer, the cryo-fixed cells as well as the supporting film layer face various mechanical forces and thermal stresses due to temperature fluctuations. Moreover, after cells are cryo-FIB milled, the resulting thin lamellae continue to endure external forces from mechanical handling and thermal stress. We propose a simple, yet highly effective modification to the standard rectangular milling pattern by implementing “fillets” or corner smoothing providing better mechanical stability. This adjustment helps to avoid sharp corners at the lamella edges, thereby reducing stress concentration. As a result, this modification decreases the likelihood of lamella breakage and improves the overall yield of ready-for-TEM lamellae by 33% as verified experimentally.

---

Cryogenic-focused ion beam (cryo-FIB) milling is a well-established technique for producing electron-transparent samples for cryo-electron tomography (cryo-ET). Typically, cells are deposited or grown on electron microscopy (EM) grids and thin cryo-lamellae are prepared by cryo-FIB milling on the plunge-frozen grid [1, 2, 3, 4, 5, 6]. These thin cellular cryo-lamellae (typically *<*200 nm thick), are then transferred into a cryo-transmission electron microscope (cryo-TEM). This allows for the capture of high-resolution images of cells in their near-native state.

After freezing and cryo-FIB milling, cryo-lamellae are exposed to stress from a number of factors, including mechanical handling and thermal cycling stress. During cryo-transfer from cryo-FIB to cryo-TEM, the lamellae can experience temperature fluctuations of several tens of degrees. These temperature variations, combined with the mismatch in thermal expansion coefficients between the grid material (typically copper), the thin carbon or gold film, and the primarily water-based cell, generate significant thermally induced mechanical stress. This stress, along with mechanical forces from grid handling, contributes to structural instability and deformations that are particularly detrimental to the delicate cryo-lamellae. As a result, the combined effects of mechanical and thermo-mechanical stresses can often lead to lamella breakage.

Cryo-FIB milling is a low-throughput serial process, and the number of lamellae produced remains relatively low. Even with automation, a skilled operator can typically produce only 15–25 lamellae in a single cryo-FIB session. Each lamella requires about half an hour of instrument time (in addition to the extensive time needed for sample preparation). Consequently, every broken lamella represents not only lost time, but also a missed opportunity to collect valuable TEM data.

The structural stability of a prepared lamella follows the same principles as any mechanical structure, whether at the microor macro-scale. When stress levels remain below a certain threshold, known as the yield point, the structure only undergoes elastic deformations and can return to its original shape. However, once stress levels exceed the yield point, the structure begins to deform irreversibly. Further increases in stress will lead to a gradual increase in deformations until the material ultimately fails or breaks.

The mechanical stability of prepared lamellae is not extensively discussed in the literature. A number of publications have suggested and/or introduced improvements to the stability and survivability of cryo-FIB-milled lamellae. Wang et al. [7] discuss the development of reliable methodologies for fabricating structurally sound large tissue lamellae and examine the challenges associated with lamellae failure (see for example Supplementary Figure 5d in [7]). Wang et al. [7] rely on disconnecting one edge of their long and wide lamella from the bulk of the sample to avoid breakage. Furthermore, they introduced a “furrow–ridge structure” strategy, characterised by alternating furrows and ribbons. The furrows represent the thinned-down areas, while the ridges are thick ribbons that enhance the mechanical stiffness of the overall structure. This approach has proven effective in maintaining lamella stability and reducing its bending. Kelly et al. have developed a “notch milling” strategy within the Waffle Method sample preparation technique [8, 9]. The notch separates one tip of the lamella by 200 nm from either side from the bulk of the sample without fully releasing it. The notch milling provides limited freedom of movement of the lamella tip which allows to dissipate some of the stress that develops in the lamella during handling and transfer. In the MEMS (micro-electromechanical systems) field, this design is known as “stoppers” or “overrange stops”, that prevent excessive structural motion due to external shocks that could lead to micro-mechanical structure failure [10, 11]. The notch milling was reported to significantly improve the Waffle-lamellae resilience and survivability during grid transfer and cryo-TEM characterisation. Wolff et al. have developed the “micro-expansion” strategy, or gap-milling on the sides of a cell lamella to improve its mechanical robustness [12]. Although the initial purpose of milling trenches on either side of the lamella was to protect it from stress and deformations in the grid that could cause its bending (also reported by Zhang et al. in Ref. [13]), this method was shown to improve the lamellae’s overall survivability. Micro-expansion gaps have become a standard feature in lamella preparation protocols and are widely adopted by the cryo-FIB community due to their simplicity, effectiveness, and ease of implementation.

Instances of broken or failed lamellae are seldom reported, with the focus predominantly on successfully prepared and aesthetically pleasing lamellae. From the limited literature reports [7], communications [14] and from our experience, the lamellae tend to crack or break close to the edges where they connect to the bulk sample. This suggests that mechanical stresses are highest at the lamellae edges. The lamellae are typically cryo-FIB-milled using rectangular patterns. The focused ion beam scans within the defined patterns above and below the region of interest and sputters away the excess biological material. Consequently, the resulting thin lamella has sharp corners at its connection edges to the bulk sample, which is where stress concentration can occur (Fig. 1, top). Here, we propose a minor but powerful modification in the standard ion beam milling practice, namely fillets on the corners of the rectangular milling patterns (Fig. 1, bottom). By switching to filleted patterns, the resulting lamellae will no longer have sharp internal corners but will rather have a smooth and rounded transition from the bulk of the sample to the lamella surfaces (Fig. 1, bottom). In structural and mechanical engineering, fillets are a standard design feature used to alleviate stress concentration at sharp corners. Fillets involve rounding the internal sharp corners of structures. By smoothing the sharp corners, fillets help redistribute the stress over a larger area, thus making the structures able to withstand a higher level of stress without mechanical failure.

**Figure 1:**
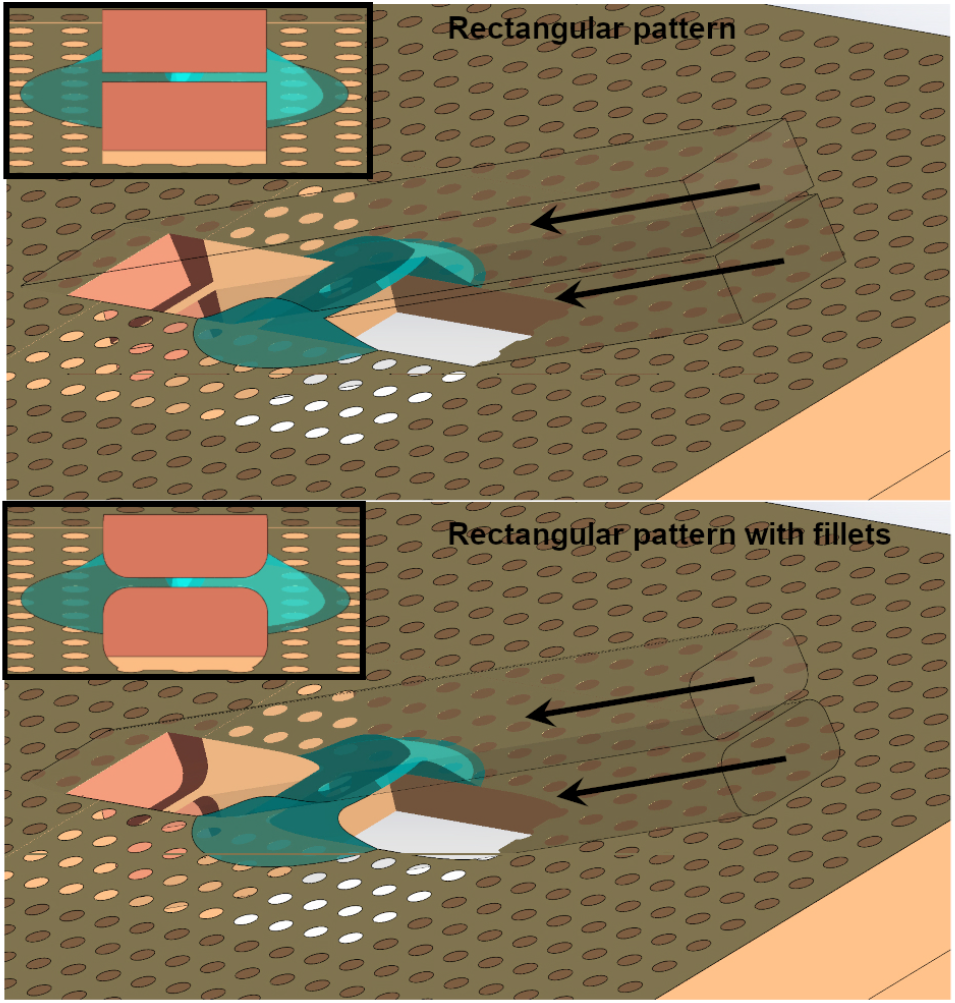
Schematic of a cell on EM-grid milled by (top) rectangular patterns and (bottom) by patterns that incorporate fillets. (top) The resulting lamella has sharp internal corners when milled with a rectangular pattern, whereas it has (bottom) smooth, rounded corners when milled with a rectangular pattern that includes fillets. The inserts show the sample viewed from the FIB column at the shallow angle (lamella-mill angle) with respect to the sample plane.

The lamella-on-grid can be analysed and treated as any other mechanical structure, with its performance under external stimuli governed by the principles of structural mechanics. This complex structure, composed of various materials such as gold, copper, carbon, water, and ice, exists in a harsh environment. It is subjected to a wide range of mechanical loads and must remain structurally sound across tens of degrees of temperature variation. Due to its complexity, the structural modeling needs to be done using numerical methods, such as Finite Element Method (FEM). Conducting a full quantitative analysis is highly challenging and demands substantial computational resources to accurately reproduce and discretise the geometry, including the complex shape of the cell. Furthermore, accurate FEM simulations require comprehensive knowledge of all temperature-dependent material properties. The accurate numerical modeling of lamella structural performance is a topic of future research. These numerical studies have the potential to further refine process parameters (for instance, the micro-expansions were not studied in simulations and the parameters can be further optimised). Nevertheless, even qualitative results from simplified simulations can support and substantiate that filleted lamellae may outperform traditional lamellae with sharp corners in terms of their survivability. Figure 2 shows qualitative finite element method (FEM) analysis conducted using the 3D CAD modeling software SolidWorks™. This analysis examines two sources of structural stress on both standard and filleted lamellae: (i) a drag force acting on the grid surface (Fig. 2a,b), simulating the grid’s motion within liquid nitrogen, and (ii) thermomechanical stress resulting from a 20-degree temperature drop (Fig. 2c,d), representing the scenario where a lamella milled at approximately 100 K is immersed in liquid nitrogen at 77 K (e.g., for storage purposes).

**Figure 2:**
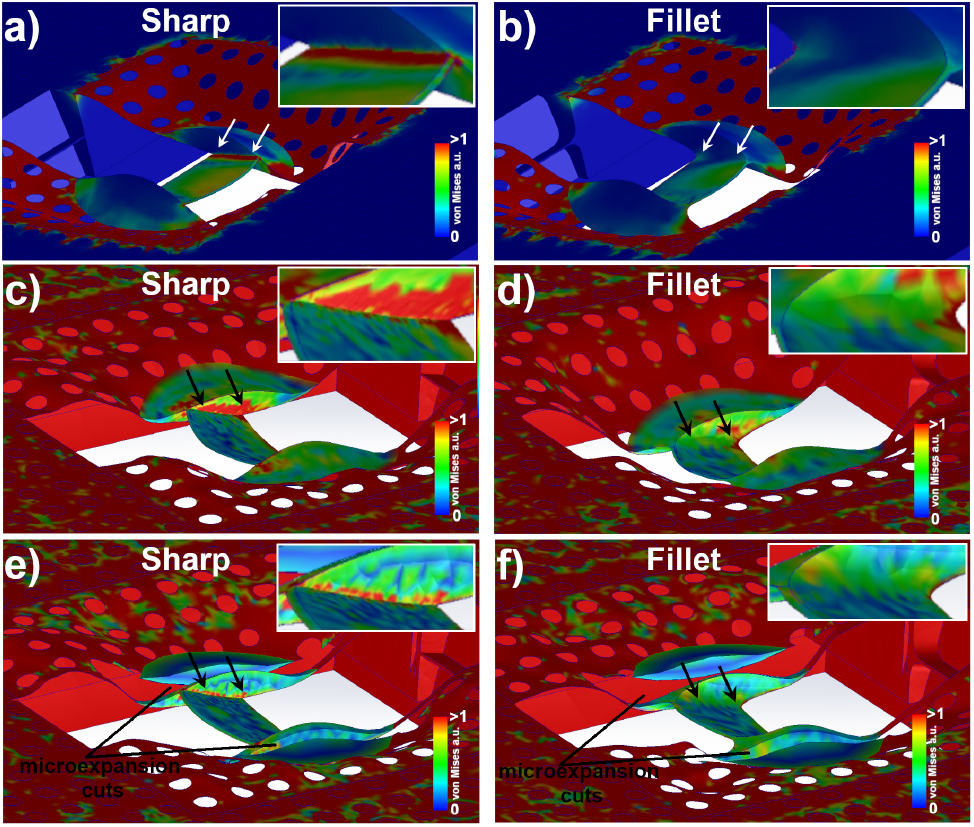
Finite Element Method (FEM) simulations of the milled lamella-on-grid mechanical response to external stimuli. (a,b) Lamella with sharp and filleted internal corners, respectively, its exaggerated deformation and von Mises stress when subjected to a drag force perpendicular to the grid surface. (c,d) Lamella with sharp and filleted internal corners, respectively, its exaggerated deformation and von Mises stress caused by the thermal stress developed due to a 20 K temperature drop. (e,f) Similar conditions to (c,d) but with microexpansion gaps milled. Von Mises stress scale is arbitrary and adjusted to highlight the stress inside the lamella (red shows stress value saturation over a threshold).

Under similar initial conditions, it is evident that stress concentration occurs around the sharp corners of the lamella (Fig. 2a). However, smoothing the corner sharpness reduces stress concentration by distributing the stress over a larger surface area (Fig. 2b). If the stress exceeds the yield strength, the lamella material may crack, leading to loss of the lamella. In contrast, the smooth-edged filleted lamella exhibits overall lower stress levels, allowing it to withstand even harsher treatment and conditions.

A lamella-on-grid sample undergoing thermal cycling, either during cryo-transfer or due to temperature fluctuations, will accumulate internal stress. This stress arises from the differences in the thermal contraction or expansion of the various materials comprising the sample, such as the copper frame, the gold or carbon holey support film, and the vitreous ice of the cell, due to their differing thermal expansion coefficients. The lamella milling process, involving portions of the sample removal and creating large openings and microexpansion gaps, facilitates the relaxation of pre-existing mechanical stresses through deformations in both the sample and the supporting carbon/gold film. Subsequently, when the sample is cooled further, the primary source of additional stress and deformation is the thermal contraction. A comparative analysis of lamella structural performance under a 20-degree temperature drop is shown in Figures 2c and d. Figure 2c shows stress concentration at a sharp internal corner (indicative of increased vulnerability at this location), while Figure 2d demonstrates a significant reduction in stress through the application of a fillet. This stress reduction enhances the lamella’s ability to withstand the imposed conditions, allowing it a greater chance to survive until the TEM characterisation.

The simulation indicates that the lamella exhibits bending and curvature following the temperature decrease. This phenomenon is attributed to the significant disparity in the thermal expansion coefficients between the ice within the cell and the underlying carbon or gold film, as well as the copper grid bars. The mechanical properties, particularly the linear thermal expansion coefficients across a broad temperature range, are well-documented and tabulated for metals [15, 16] and, to some extent, for thin metal films [17]. While the temperature-dependent properties of crystalline ice are extensively researched and tabulated [18, 19], data for vitreous ice remains limited, with only a few experimentally measured data points available [20]. This uncertainty is even greater for vitrified cells and tissue, that contain a mixture of water and organic molecules. Using the data for crystalline ice, the thermal expansion coefficient for ice is substantially higher (60×10^−6^ K^−1^ at room temperature dropping to ∼12.5×10^−6^ K^−1^ at 100 K) compared to copper (16×10^−6^ K^−1^ at room temperature dropping to ∼10×10^−6^ K^−1^ at 100 K), gold (14×10^−6^ K^−1^ at room temperature decreasing to ∼11×10^−6^ K^−1^ at 100 K), or amorphous carbon films (2–10×10^−6^ K^−1^ at room temperature [21]). The considerable discrepancy in thermal expansion between the ice and the materials comprising the grid and the supporting film results in greater contraction of the cell and lamella, leading to deformation of the supporting film and bending of the thin lamella. Figures 2(e,f) illustrate the effects of implementing microexpansion cuts in a lamella, both with and without inner corner fillets, under thermal stress conditions. The microexpansion gaps effectively shield the lamella from some of the stresses and deformations in the surrounding grid film and the bulk of the cell. However, while these gaps reduce the mechanical stress levels in the lamella compared to scenarios without them, they are ineffective against the stresses and deformations that develop within the lamella due to temperature variations or mechanical loads. Therefore, employing both strategies – microexpansion gaps and corner fillets – in tandem is highly advantageous. This combined approach enhances the robustness of the lamellae, improves their resilience during cryo-transfers, and increases their survivability, ultimately leading to higher data collection yields.

Corner fillets and chamfers are standard features in CAD design software (e.g., AutoCAD™ or SolidWorks™). However, to the best of our knowledge, no cryo-FIB vendor currently offers a fillet-rectangle as a common milling pattern in their library of available beam scanning and milling geometries. To experimentally verify the improvement in mechanical robustness of lamellae with fillets, we used the highly customisable open-source Autolamella v2 application, that is based on the OpenFIBSEM API for microscope control [22, 23]. We were able to introduce the fillets in patterns by combining standard rectangular and circular shapes (Fig. 3). Here, the fillet pattern is characterised by a single parameter - the radius of the fillet. It would be straightforward for cryo-FIB vendors to introduce a fillet into rectangular and cleaning cross-section patterns typically used in lamella milling.

**Figure 3:**
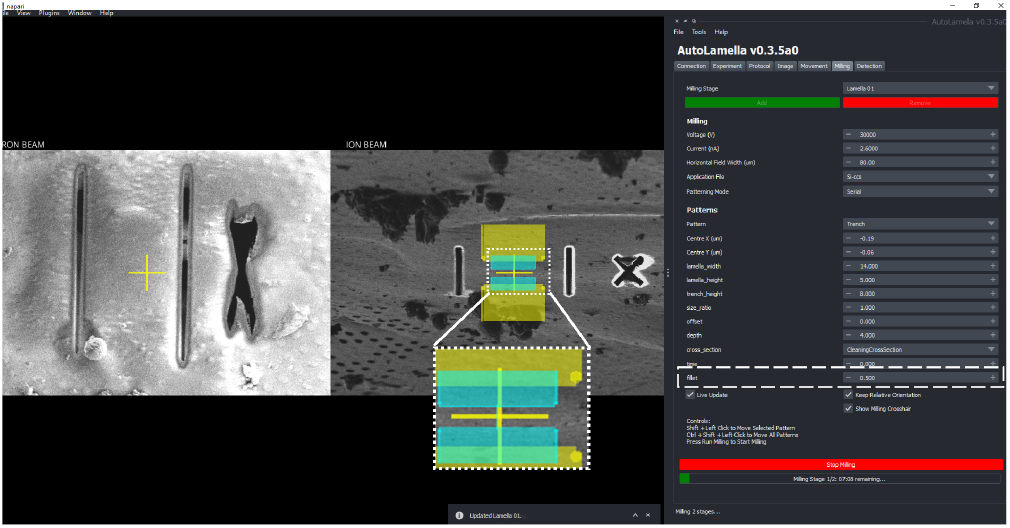
Autolamella main GUI window showing progress of a lamella milling. A fiducial marker and the microexpansion gaps have been milled (right - scanning electron beam view, left - corresponding view in the focused ion beam view). The rough trench milling patterns are shown in yellow, while the intermediate milling patterns are shown in cyan. Autolamella is highly customisable. Here we introduced fillets built from basic available patterns such as rectangles and circles using a single parameter “fillet radius” as shown in the GUI pattern control.

Fig. 4 compares prepared sharp-edge lamella (a-c) with a lamella incorporating fillets (d-f) of different radii for different milling stages (500 nm, 200 nm and 100 nm, for rough, intermediate and fine polishing steps, respectively.). To assess the anticipated enhancement in lamella robustness resulting from the incorporation of fillets, we milled over 150 lamellae with and without fillets. Table 1 summarises the results. Among the 150 conventional lamellae and 156 fillet-lamellae, 112 and 130, respectively, were of high quality. These lamellae were fully released at both ends, appropriately thinned for cryo-ET experiments, were flat, uniform and undeformed, and contained biologically relevant structures. The remaining 38 conventional and 26 fillet-lamellae were of lower quality, being either devoid of biological structures, deformed, minorly nonuniform, irregularly milled, or containing curtains. Nevertheless, we include these lower quality lamellae in the overall statistics, as our primary concern is the survivability of milled structures rather than their biological relevance. Among the structurally sound lamellae, 62% of the conventional lamellae and 82% of the fillet-lamellae successfully survived the cryo-transfer from the cryo-FIBSEM to the cryo-TEM, where they could be observed. This represents an improvement of nearly 33% (one-third) in the survival rate of fillet-lamellae during transfers compared to the conventional structures, ensuring their intact condition for subsequent TEM studies. Although lamella robustness is influenced by numerous factors, including the type of grids used, the type of cells seeded onto the grids, the plunge freezing procedure, general grid handling, and other variables, this study ensured that the grids were treated uniformly, lamellae had identical or comparable dimensions and had uniform thickness (160–180 nm). The primary variable was the presence of fillet corners, enabling a systematic comparison and evaluation of their impact on the overall structural health of the lamellae.

**Table 1:** Summary of the lamella survivability for standard lamellae (sharp corners) and lamellae with fillets (rounded corners). Both types of lamellae were prepared with microexpansion gaps.

| Lamellae | Suitable<br>for TEM | (Good<br>quality) | Intact<br>to TEM | Intact<br>% |
| --- | --- | --- | --- | --- |
| Standard | 150 | (112) | 93 | 62% |
| Fillets | 156 | (130) | 128 | 82% |

**Figure 4:**
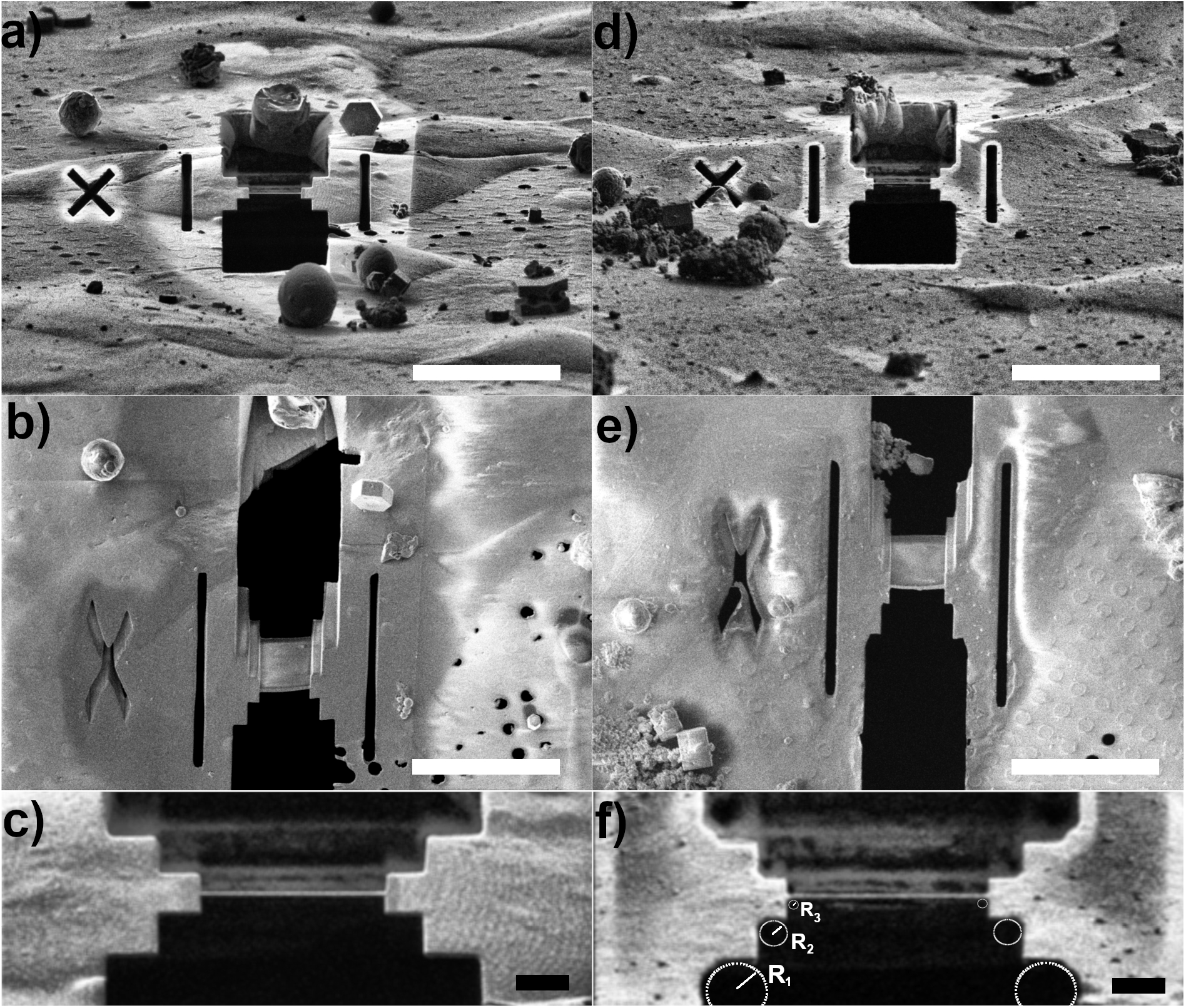
Micrographs of MEF cell lamellae prepared without (a,b,c) and with (d,e,f) the inner corner fillet. (a,d) Focused ion beam view (14^°^ ion beam to the grid surface). (b,e) Corresponding electron beam view of the prepared lamellae. (c,f) Magnified view of (a,d) comparing the sharp inner corner with the filleted inner corner. Scale bar in (a,b,d,e) is 20 *µ*m, and 2 *µ*m in (c,f).

Figure 5 shows TEM characterisation of a lamella with corner fillets. Despite the broken support film at the right-hand microexpansion gap, the lamella successfully survived cryo-transfers from the cryo-FIB to the cryo-TEM. The breakage occurred due to the pre-existing cracks in the support film prior to milling. Nonetheless, the lamella remained structurally sound and flat, even with reduced support from the underlying grid. Once it was inside the TEM, a crack developed at the centre of the lamella, whereas cracking typically occurs at the edges. This crack likely emerged due to the lamella’s ability to twist around its axis due to the absence of a fixed support, and due to the thickness non-uniformity at its base. Ordinarily, such a lamella would be prone to breakage and total loss; however, this lamella not only survived significant structural failures but was still usable for high-quality TEM tomogram data collection (Fig. 5d).

**Figure 5:**
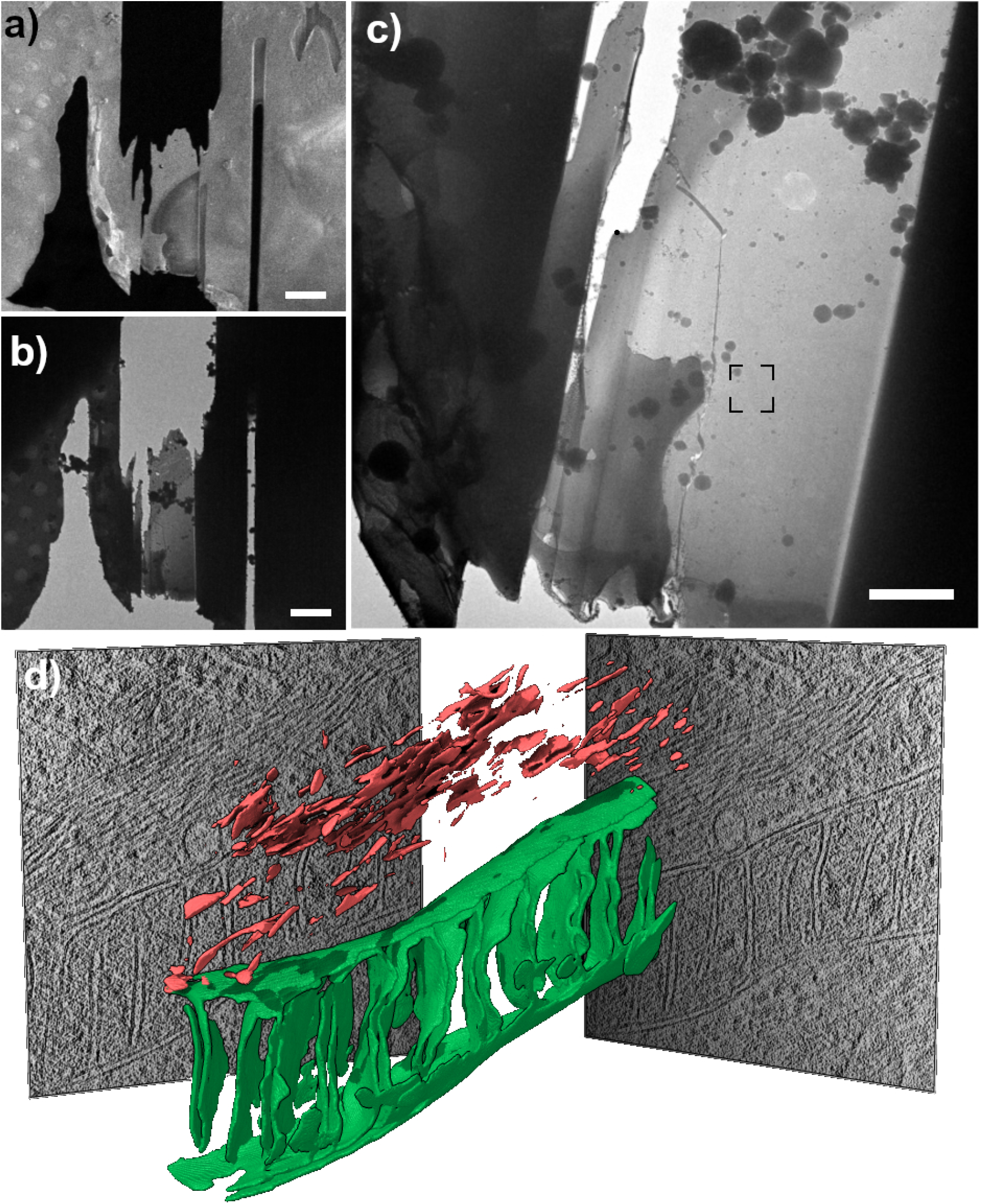
Cryo-electron tomography of a NCI-H441 cell lamella milled with corner fillets. a) SEM micrograph of a prepared lamella. b) The same lamella imaged in a 300-keV cryo-TEM at very low magnification (740*×*) after the cryo-transfer from the cryo-FIB into the cryo-TEM. c) Low magnification TEM image (3600*×*) of the lamella in b). d) Segmentation of a mitochondrion (green) and filaments (red) from the tilt-series collected on the selected area in (c) and projection images of the centre of the 3D reconstructed volume. The field of view of the reconstruction is 1.6*×*1.4 *µ*m. Scale bar in (a,b) is 5 *µ*m, 2 *µ*m in (c).

In summary, we demonstrate an enhancement in the yield of lamella preparation using cryo-FIB milling. The implementation of microexpansion gaps improves lamella yields by shielding the lamellae from stresses and deformations in the grid support. Here, we introduce fillets on the inner corners of the lamellae—rounded corners that help reduce stress concentration—which protect the lamellae from stresses and deformations that accumulate within the lamella itself. The introduction of fillets has improved the yield by one-third. Although the incorporation of fillets represents a minor modification to existing cryo-FIB lamella preparation protocols, it significantly enhances sample throughput and facilitates the collection of more TEM data. This study represents the first attempt at mechanical and structural simulations of lamella performance, and we believe that further optimisation of lamella preparation parameters can enhance lamella survivability in the harsh environments to which they are subjected.

## Materials and methods

### Cell culture on EM grids and freezing

Two types of cells were cultured for this study. Firstly, mouse embryonic fibroblasts (MEF) were maintained at 37 °C, 5% CO_2_ in in-house made DMEM High Glucose (DMEM) supplemented with 10% FBS and GlutaMAX™ Supplement (Gibco #35050061). The cells were then seeded onto EM grids (Quantifoil Au R2/2) after coating the grids with ∼3 nm carbon using a Leica AF200 (Leica Systems), and glow-discharging for 30 s using a Pelco EasyGlow™. After 48 hours, the cells grown on the grids were transferred from the culture room to the EM facility. Freezing conditions were set up in a Leica EM-GP2 (liquid ethane held at -175 °C, chamber at 37 °C and 95% relative humidity). Samples were backside-blotted for 12 s and plunge-frozen into liquid ethane. After plunging, the grids were placed into a grid box kept in liquid nitrogen to prevent devitrification and limit ice contamination.

Secondly, NCI-H441 human lung cancer cells were cultured in RPMI 1640 media supplemented with 10% fetal bovine serum (FBS) and 1% penicillin-streptomycin in a 37 °C incubator with 5% CO_2_. The cells were deposited onto glow-discharged EM grids (Quantifoil Au R2/2), and backside-blotted for 6 to 8 s in a Leica EM-GP2 using freezing conditions similar to those used for the MEF cell line.

### Cryo-FIB milling

Lamellae fabrication was performed on a dual-beam ThermoFisher Scientific (TFS) Helios 5 UX Cryo-FIBSEM using 30 keV Ga+ beam. The grids clipped in autogrids were cryogenically transferred into the cryo-FIB, and suitable cells for thinning and milling were selected from mapping and imaging the grids with electron and ion beams. The ion beam incidence with respect to the grid plane was set to 12^°^ or 14^°^ and kept constant throughout the lamella milling. We ensured that each selected milling site was at the eucentric point of the microscope, e.g., the point of coincidence between the electron and ion beams. In this point the sample is at 4 mm from the electron microscope’s polepiece. Next, the sample was lowered to 5.5 mm from the electron polepiece and a layer of organometallic platinum was deposited onto the grids using gas injection system (GIS) operated at 29 °C for 12 s. The lamellae milling was aided with Autolamella v2 software [22, 23]. First, we translated the stage from one lamella site to the next, milling fiducial markers at each site to facilitate future alignments. Next, we proceeded to each lamella site, aligning using the previously milled fiducial markers. We then performed rough and intermediate milling with beam currents of 2.6 nA and 0.44 nA, respectively. Before the rough milling, we created micro-expansion gaps using a 2.6 nA beam current. The rough milling step aimed at creating a 5-*µ*m-thick and 14-*µ*m-wide lamella, while the intermediate milling step aimed at creating a 1.5-*µ*m-thick and 10-*µ*m-wide lamella. The microexpansion gaps were positioned on either side of the lamella 10 *µ*m from its centre. The polishing of all thick lamellae was performed sequentially using either a 41 pA or 90 pA beam current to produce lamellae thinner than 200 nm. We moved from one lamella site to another, realigned the position using the previously milled fiducial markers, and conducted the final polishing. The Autolamella v2 has recorded the process and intermediate steps in images. All SEM images were collected at 2 keV beam energy and 0.1 nA beam current. All FIB images were acquired at 30 keV beam energy and at 41 pA beam current.

### Cryo-TEM tomography

After FIB milling, EM grids were promptly (within 12 hours) transferred into a Titan Krios G4 (ThermoFischer Scientific) cryo-TEM, operating at 300 kV and equipped with a Selectris-X energy filter and a Falcon 4i direct electron camera. Very-low magnification (740×) and low magnification (3,600×) images of the cryo-FIB lamellae present in this work were collected as part of the tomography workflow using the TFS Tomography TOMO5 software. Electron-Event Representation (EER) tilt-series were collected at 64,000× magnification (corresponding pixel size 1.97 Å) using a 20 eV slit. Images were collected between -42^°^ and +70^°^ stage tilts every 2^°^ using a dose-symmetric scheme [24], corresponding to a ±56^°^ rotation around the angle at which the lamellae were milled (about +14…+15^°^). The total electron fluence used per tilt-series was 100-120 e^−^/Å^2^.

### Cryo-TEM alignment, reconstruction and segmentation

EER images were aligned and averaged with MotionCor3 using 4-by-4 patches [25], outputting even/odd image pairs as well, for subsequent use of CryoCARE [26, 27]. The tilt-images containing all frames were sorted to generate the tilt-ordered raw tilt-series using a custom script and aligned with AreTomo2 [28] using a matrix of 4-by-4 patches. The AreTomo2 alignment was imposed on the even/odd tilt-series pairs and twin 3D reconstructions binned 4× (pixel size 7.88 Å) were computed in Tomo3D using Tegunov’s filter [29, 30]. Denoising was performed in CryoCARE using standard parameters. The denoised 3D reconstruction was binned another time (resulting pixel size 15.76 Å) and then processed with IsoNet (deconvolution and missing-wedge filling) [31]. Automatic segmentation of the resulting corrected 3D reconstruction was performed using membrain-seg [32], manually corrected, and rendered in ChimeraX [33].

## Acknowledgement

The authors acknowledge the use of instruments and assistance at the Monash Ramaciotti Centre for Cryo-Electron Microscopy, a Node of Microscopy Australia. This research used equipment funded by Australian Research Council grants: FEI Helios Cryo FIBSEM - ARC LIEF (LE150100132) and Titan Krios - ARC LIEF (LE120100090). This research was in part funded by ARC DP200103637. This work has been made possible in part by CZI grant DAF2021-225399 and grant DOI https://doi.org/10.37921/334038myxhsa from the Chan Zuckerberg Initiative DAF, an advised fund of Silicon Valley Community Foundation (funder DOI: 10.13039/100014989). This research was supported by the Australian Research Council Industrial Transformation Training Centre for Cryo-Electron Microscopy of Membrane Proteins for Drug Discovery (IC200100052) and funded by the Australian Government. LY and VN are grateful to Associate Professor Daniel Gough at Hudson Institute of Medical Research for providing NCI-H441 cells for this study.

